# Varying perivascular astroglial endfoot dimensions along the vascular tree maintain perivascular-interstitial flux through the cortical mantle

**DOI:** 10.1101/2020.07.15.204545

**Authors:** Marie Xun Wang, Lori Ray, Kenji F. Tanaka, Jeffrey J. Iliff, Jeffrey Heys

## Abstract

The glymphatic system is a recently defined brain-wide network of perivascular spaces along which cerebrospinal fluid (CSF) and interstitial solutes exchange. Astrocyte endfeet encircling the perivascular space form a physical barrier in between these two compartments, and fluid and solutes that are not taken up by astrocytes move out of the perivascular space through the junctions in between astrocyte endfeet. However, little is known about the anatomical structure and the physiological roles of the astrocyte endfeet in regulating the local perivascular exchange. Here, visualizing astrocyte endfoot-endfoot junctions with immunofluorescent labeling against the protein megalencephalic leukoencephalopathy with subcortical cysts-1 (MLC1), we characterized endfoot dimensions along the mouse cerebrovascular tree. We observed marked heterogeneity in endfoot dimensions along vessels of different sizes, and of different types. Specifically, endfoot size was positively correlated with the vessel diameters, with large vessel segments surrounded by large endfeet and small vessel segments surrounded by small endfeet. This association was most pronounced along arterial, rather than venous segments. Computational modeling simulating vascular trees with uniform or varying endfeet dimensions demonstrates that varying endfoot dimensions maintain near constant perivascular-interstitial flux despite correspondingly declining perivascular pressures along the cerebrovascular tree through the cortical depth. These results describe a novel anatomical feature of perivascular astroglial endfeet and suggest that endfoot heterogeneity may be an evolutionary adaptation to maintain perivascular CSF-interstitial fluid exchange through deep brain structures.

## Introduction

The brain is unique among organs in that the exchange of free solutes between the interstitial compartment and blood is restricted by the brain blood barrier (BBB)^1^, such that solutes that are not degraded locally and lack a specific BBB efflux transporter must be cleared through exchange with the cerebrospinal fluid (CSF) compartment. The CSF is a relatively low-protein and cell-free fluid secreted by the choroid plexus that circulates through the ventricular system to fill the subarachnoid space and cisternal compartments^2^. CSF also extends from the subarachnoid space into brain tissue along perivascular spaces, or Virchow-Robin spaces, surrounding penetrating cerebral blood vessels^3^. These are fluid-filled annular spaces between the vessel wall of penetrating vessels and specialized perivascular astrocyte endfeet surrounding them^3^. Perivascular astroglial endfeet cover nearly the entire surface area of the cerebral vasculature, and play important role in maintaining the integrity of the BBB and in regulating cerebral blood flow^4–7^. Unlike the vascular endothelial cells, the junctions between overlapping astrocyte endfeet lack tight junctions, which permits the movement of fluid and solutes from perivascular spaces to exchange with the wider brain interstitium^8^.

Although fluid movement along perivascular spaces has been observed for decades^9–11^, much recent interest has followed the description of the ‘glymphatic’ system, a brain-wide network of perivascular spaces along which CSF enters and interstitial solutes are cleared from brain tissue^12,13^. These recent studies have demonstrated that perivascular exchange is driven in part by vessel pulsation^14,15^, is supported by the perivascular astroglial water channel aquaporin-4 (AQP4)^12,16^, is more rapid in the sleeping compared to the waking brain^17,18^. Studies in rodents and in human subjects suggest that impairment of glymphatic exchange contributes to a wide range of neurological conditions including Alzheimer’s disease^19^, stroke^20^, cerebral small vessel disease^21^, traumatic brain injury^22^, subarachnoid hemorrhage^23^ and migraine^24^. Therefore, it is crucial to understand the anatomical basis, molecular underpinnings and physiological regulation of perivascular glymphatic exchange.

AQP4 is an astroglial water channel that localizes primarily to the perivascular endfeet through its association with the dystrophin-associated complex (DAC)^25^. Weighted Gene Coexpression Network Analysis (WGCNA) from two publicly-available human brain transcriptomic datasets demonstrated that the expression of *AQP4* and other DAC-associated genes was associated with expression of the *MLC1* (megalencephalic leukoencephalopathy with subcortical cysts-1)^26,27^, an astrocyte-specific gene of unknown function but with limited homology (<20% protein identity) to the Kv1.1 potassium channel. Mutations in the *MLC1* gene cause profound white matter degeneration^28,29^. Evaluation of *MLC1* expression in a human dementia dataset revealed that changes in MLC1 expression were associated with cortical tau pathology and clinical dementia^27^.

The physiological function of MLC1 remains unknow. Previous work done in cell culture in *Mlc1*-null mice suggest that MLC1 may be involved in astrocyte ion and volume regulation through its interaction with different chloride, potassium and cation channels, or with gap junction-forming connexin-43^30–35^. Anatomically, immunogold-electron microscopic images in postmortem human tissue showed that MLC1 localized to the junctions connecting overlapping perivascular astrocytic endfeet^36^. In this study, we took advantage of the MLC1 expression at the astrocyte endfoot-endfoot junctions to describe the patterns of astrocyte endfoot ensheathment along the cerebrovascular tree. Observing heterogeneity of endfoot size along cerebral blood vessels, we employed a computational model to describe the role that this variation may play in maintaining exchange between perivascular spaces and the brain interstitium through the depth of brain tissue and branching order.

## Results

### MLC1 localization along large blood vessels

MLC1 and AQP4 expression and localization along cortical and hippocampal blood vessels were defined by immunofluorescence double-labeling in 3, 12 and 15 months old C57Bl/6 mice. As expected, AQP4 immunofluorescence was restricted predominantly to perivascular endfeet surrounding cortical and hippocampal vessels (**Figure 1A**), including along both capillaries (white arrowheads) and along larger intraparenchymal vessels (yellow arrows). While MLC1 localization was similarly perivascular, it was restricted primarily to perivascular endfeet surrounding larger intraparenchymal vessels, rather than capillaries. Using super resolution structured illumination microscopy (SR-SIM), we further characterized the disposition of MLC1 in perivascular endfeet surrounding intraparenchymal vessels. MLC1 immunofluorescence formed a ‘meshwork’ surrounding blood vessels (**Figure 1B**), in line with prior studies demonstrating with electron microscopy that MLC1 localizes to astrocyte-astrocyte junctions[cite]. MLC1 did not directly co-localize with AQP4, suggesting that while AQP4 appears to occupy most of the perivascular astroglial endfoot surface area abutting the cerebral vasculature, MLC1 marks the boundaries of overlapping perivascular endfeet surrounding non-capillary vessels in both the cortex and hippocampus.

**Figure 1:**
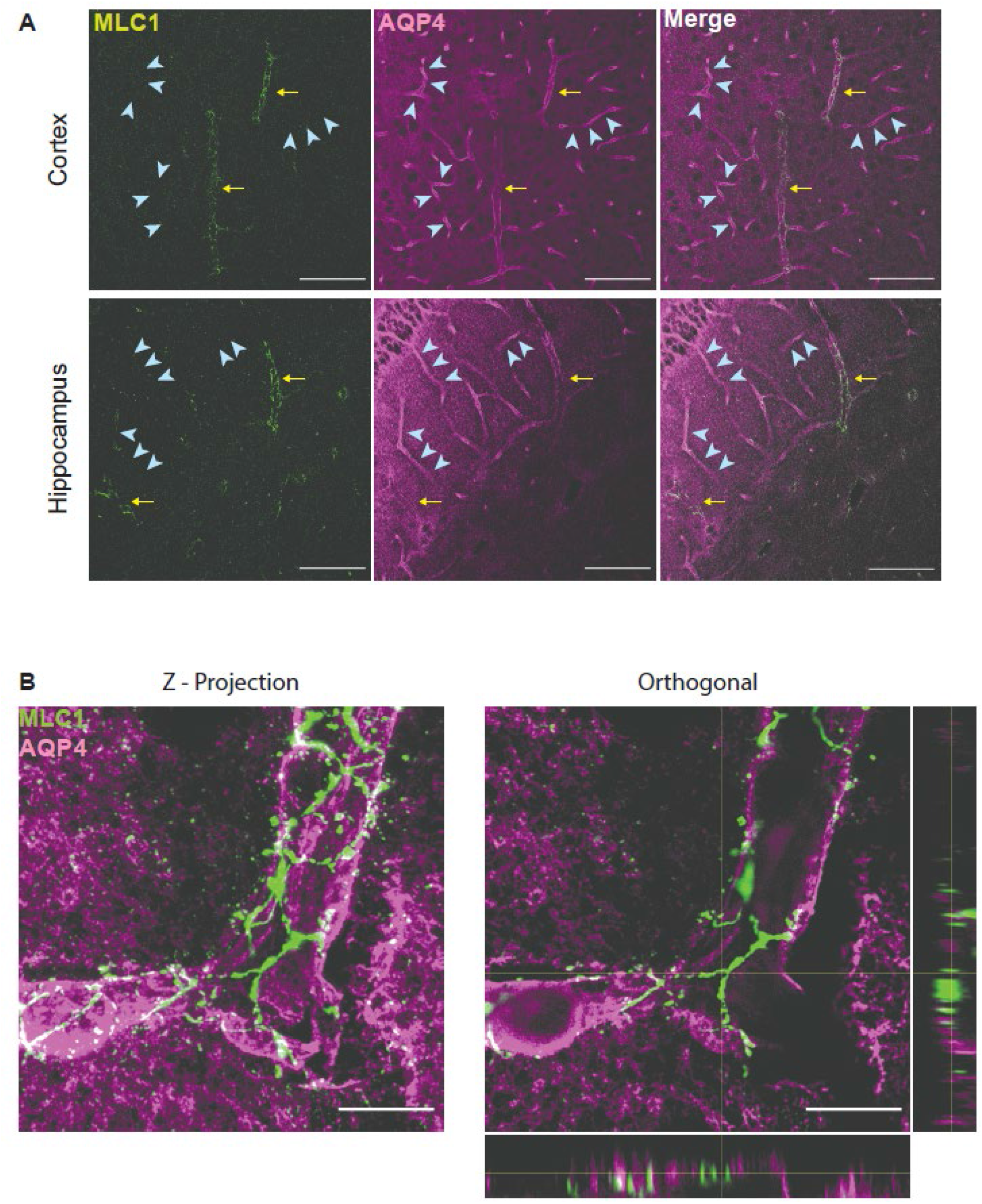
MLC1 is a marker for astrocyte endfoot-endfoot junctions along non-capillary vessels. **(A)** Confocal images of the cortex and hippocampus of 3-month old wildtype mice were taken at 20x magnification. AQP4 was used to label perivascular endfeet surrounding all blood vessels. MLC1 expression aligned with non-capillary vessels (yellow arrows) but not capillaries (blue arrow heads). Scale bar = 100 *μ*m. **(B)** MLC1 labeling defined the outline of astrocyte endfoot-endfoot junctions. Super Resolution Structured Illumination Microscopy (SRSIM) images of cortical vessels of 3-month wildtype mice were taken at 100x magnification. Maximum Z-projection (left) and a single XY plane with orthogonal XZ and YZ projections (right) suggest that MLC1 doesn’t colocalize with perivascular AQP4, but rather localizes to the perivascular endfoot-endfoot junctions. Scale bar = 10 *μ*m.

### Perivascular endfoot size declines with vessel diameter

The localization of MLC1 to perivascular endfeet surrounding intraparenchymal vessels permitted the dimension of these endfeet to be evaluated along the cerebrovascular tree. We next used whole-slice fluorescence imaging of MLC1 immunofluorescence to characterize perivascular endfoot dimensions across vessels from 3- and 15-month old animals. Initial inspection of MLC1 immunofluorescence revealed variable perivascular endfoot sizes along and between blood vessels (**Figure 2A**). Individual endfeet were manually outlined, and vessel diameters, endfoot areas and perimeters were measured (**Figure 2B**) from 3 forebrain sections per animal, 6 animals per age. Plotting the areas of all measured endfeet as a function of corresponding vessel diameter, we observed a significant positive association between vessel diameter and endfoot area (**Figure 2C**, P<0.0001, R^2^=0.1840 for pooled 3- and 12-month old animals). The relationship between vessel diameter and endfoot size did not significantly differ between 3- and 12-month old animals (P=0.3695). When endfoot values along individual vessels were averaged, the same association between increasing vessel diameter and increasing perivascular endfoot area was observed (**Figure 2D**, P<0.0001, R^2^=0.5651 for pooled 3- and 12-month old animals). Similar results were observed when median endfoot areas were calculated for each vessel (**Figure 2E**, P<0.0001, R^2^=0.4672 for pooled 3- and 12-month old animals). The relationship these parameters did not differ between 3- and 12-month old animals (P_mean_=0.1328; P_median_=0.1167). These findings demonstrate that perivascular astroglial endfeet are not uniform in size along the cerebral vasculature, but rather that as vessels decline in diameter, the area of the perivascular endfeet surrounding them decline proportionally.

**Figure 2:**
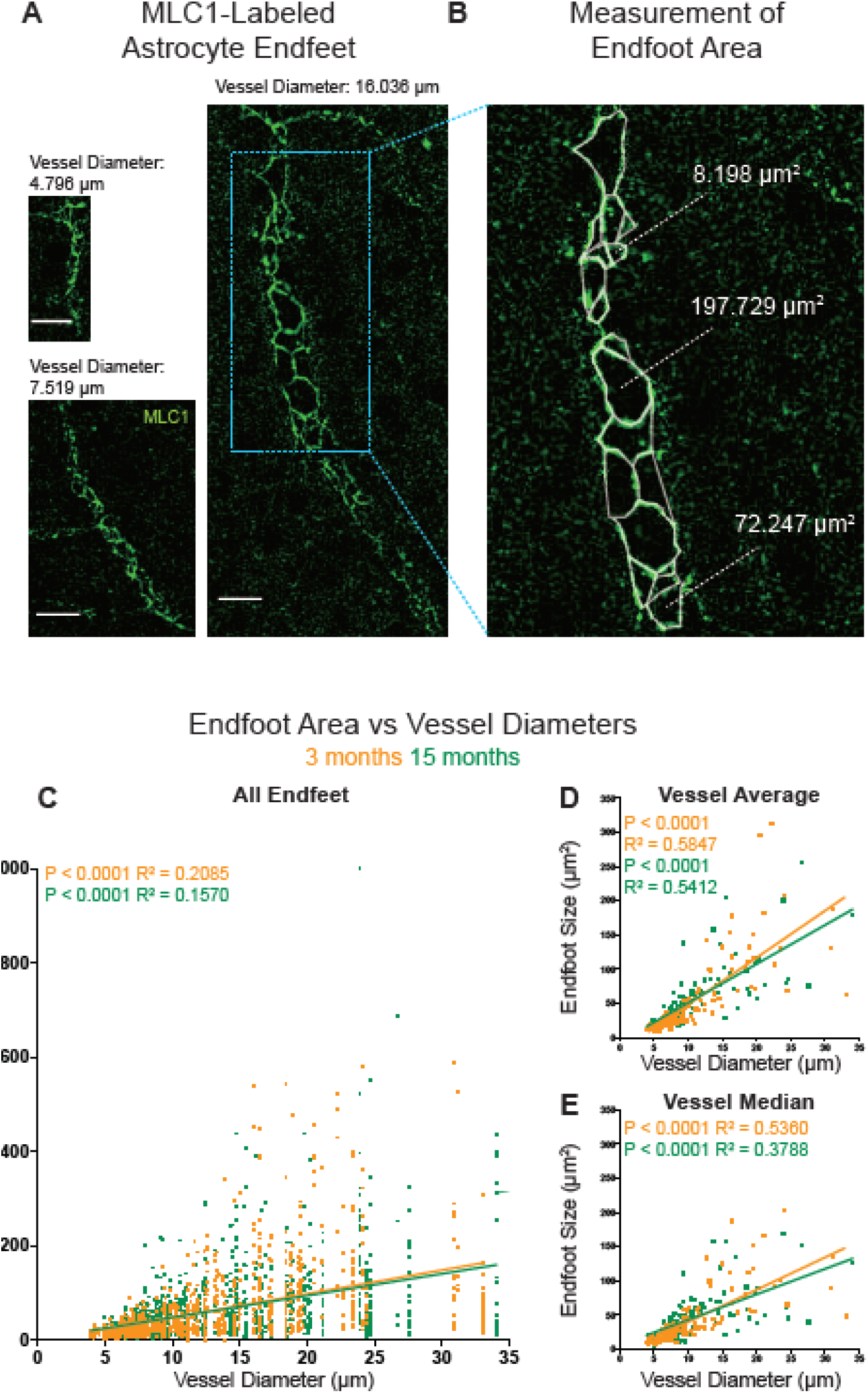
Perivascular endfoot size increases with vessel diameter and does not differ with age. **(A)** Representative confocal images show the heterogeneity of endfoot sizes along vessels of variable diameters. Larger vessels were surrounded by larger endfeet and smaller vessels were surrounded by smaller endfeet. Scale bar of all three images = 20 *μ*m. **(B)** Endfoot areas were measured by manually outlining individual endfoot domains. Vessel diameter was also measured for each vessel. **(C)** Endfeet sizes and vessel diameters of the cortical vessels of 3- and 15-month old wildtype mice were collected (3 slides per mice, 6 mice per age group). Individual endfoot areas were plotted against vessel diameters using linear regression model. A significant positive association was observed between vessel diameter and endfoot area (P_3-month_ < 0.0001, R_3-month_^2^ = 0.2085, Slope_3-month_ = 5.025. P_15-month_ < 0.0001, R_15-month_^2^ = 0.1570, Slope_15-month_ = 4.626) and no significant difference was observed between two ages. (P = 0.3695). **(D)** Similar analyses between vessel diameter and average endfoot size for each vessel and **(E)** vessel diameter and median endfoot size for each vessel showed the same results.

### Modeling the effect of endfoot size on perivascular exchange

Contrast-based studies carried out in rodents^13^, non-human primates^23^, and human subjects^37–39^ demonstrate CSF from cisternal compartments travels along perivascular spaces surrounding leptomeningeal vessels and enters the brain along perivascular spaces surrounding penetrating vessels (also termed Virchow-Robin spaces). From here, solutes such as fluorescent tracers or gadolinium-based MRI contrast agents that do not cross the blood brain barrier and are not taken up by astrocytes exchange into the brain interstitium via gaps between overlapping perivascular astroglial endfoot processes. We next utilized a computational modeling approach to explore the functional implications of the observed non-uniform perivascular astroglial endfoot dimensions on solute exchange and fluid flow between perivascular spaces and the surrounding interstitium.

Fluid flow in the perivascular space was described using an adapted version of the model described by Faghih and Sharp^40^ and includes vessels with 11 branching orders spanning vessel diameters from 1-5% larger than the largest measured vessel diameter down to the those of capillary diameter (~5 μm). The radius of daughter branches across all orders were approximated using Murray’s law of bifurcations for symmetric trees, and the length of each vessel was approximated as 20-times its radius. Flow in the annular perivascular space was assumed to be laminar, and the relationship between flow rate and pressure drop in the perivascular space followed Faghih and Sharp^40^. The total flow rate of fluid in the perivascular space was assumed to be 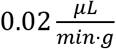 (approximated from ^10,41^), but the overall results are normalized to be independent of this value because it is difficult to accurately measure. The computational model predicted an approximately linear decrease in the pressure within the perivascular space for increasing branch order if a limited number (1 to ~15) of branching orders are considered^42^ (**Figure 3B**, grey triangles).

**Figure 3:**
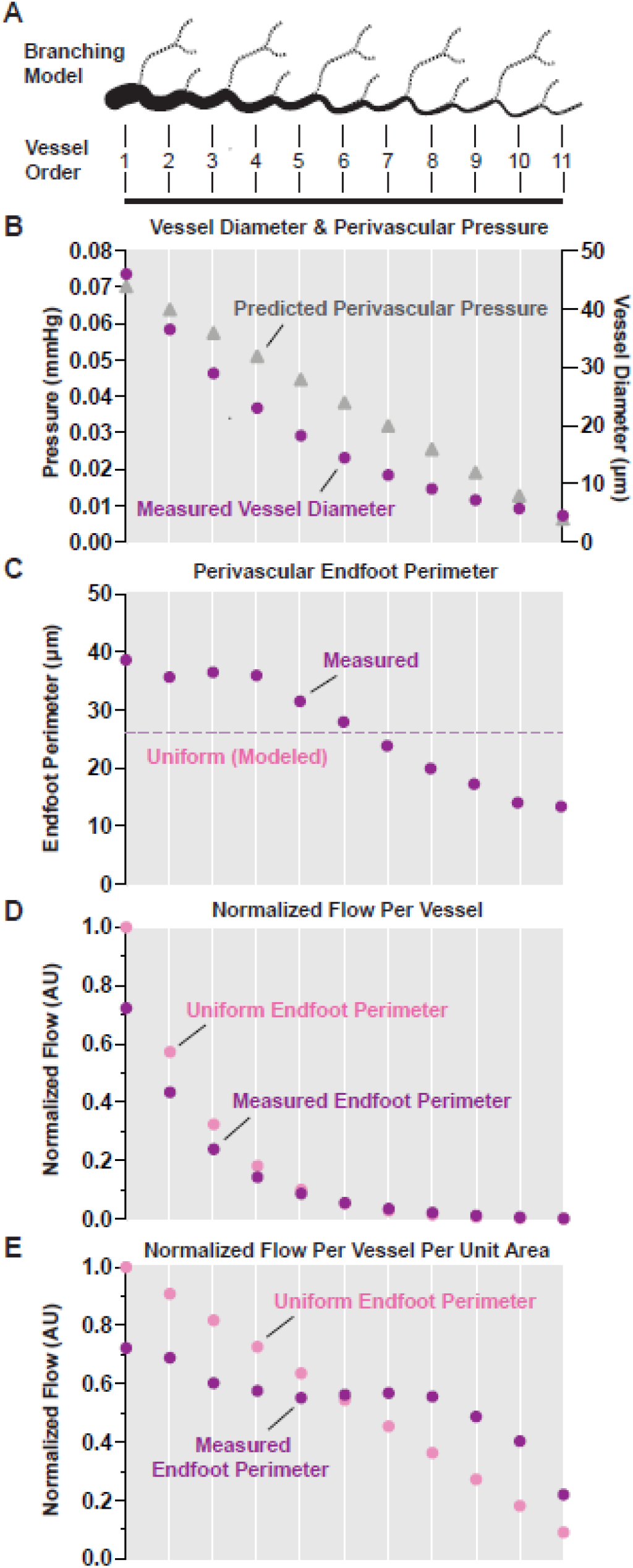
Computational modeling: Changing endfoot dimentsions lead to more uniform perivascular flux along the vascular tree. **(A)** A computational model of an 11-branching order vascular tree was generated. **(B)** Predicted perivascular pressure declined approximately linearly with increasing vessel branch order (gray triangles). Average measured vessel diameters mapped to the computational model are represented by purple circles. **(C)** The average measured perivascular endfoot perimeter for each vessel branch order was plotted (purple circles) and showed declining endfoot dimensions between 3rd and 10^th^ order branches. An overall average endfoot perimeter was calculated from all measurements (pink dashed line) and used to model the effect of constant endfoot dimensions along the model vascular tree. **(D)** When endfoot dimensions are held constant (pink circles), normalized flow from perivascular space into the interstitium declines with increasing vessel order as perivascular pressure declines. Modeling with measured endfoot dimensions that vary with branching order (purple circles) reduced predicted flux along larger vessel segments, and increased flux along small-diameter segments. **(E)** Normalized flux (flow per unit area) declined approximately linearly when endfoot dimensions were held constant (pink circle). Modeling with measured endfoot dimensions showed that varying endfoot sizes with vessel order (purple circles) reduced flux along large diameter segments, increased flux along small segments, and maintained flux largely constant between 3^rd^ and 9^th^ order segments.

Mapping the endfoot geometry data onto the idealized vascular network begins by mapping each endfoot in the data to the appropriate branching order in the model network based on the corresponding vessel diameter. **Figure 3B** (purple circles) shows mean measured vessel diameter for vessel orders 1-11. The average endfoot perimeter was estimated by assuming an approximately hexagonal endfoot shape (normalized results are not sensitive to this shape assumption based on testing of both hexagonal and pentagonal shapes). **Figure 3C** (purple circles) shows mean measured endfoot circumferences for each branching order, as well as the mean circumference across vessel segments of all diameters (pink dashed line). As noted above (**Figure 2**), the perivascular space of the largest vessels was surrounded by endfeet of larger surface area and perimeter, while small endfeet surrounded the spaces of smaller diameter vessels.

For each branching order in the model network, the total number of endfeet required to cover the perivascular space was calculated using both: (1) the average endfoot size for each branching order, and (2) the overall average endfoot size for all vessels. Based on the number of endfeet required to cover the perivascular space of the model vessels, the perimeter of the endfeet, and the predicted pressure in the model’s perivascular space, the flowrate from the perivascular space into the interstitial space was estimated at each branching order. Since the flowrate is estimated using an unknown permeability for the gap between the endfeet, the flowrate is normalized to the maximum flow rate from the largest (or root) branch, which has both the highest pressure in the perivascular space as well as the largest surface area. The normalized flowrate using measured endfoot dimensions for each branching order is shown in **Figure 3D** (purple circles), as is the normalized flowrate calculated using the endfoot diameter averaged across all branching orders (pink circles).

For the largest branches with the highest perivascular space pressures, using uniform endfoot dimensions results in up to 25% higher normalized flow rates than using endfeet dimensions (**Figure 3D**) that vary with vessel size. This is because a larger number of endfeet with an average uniform area are required to cover the larger vessels, resulting in more perimeter gaps and hence, more flow. However, if the larger branches are covered with larger area endfeet, such as in the measured values, then there are fewer perimeter gaps and less flow into the interstitial space. Although much smaller in absolute magnitude, the opposite is observed for the smallest vessels with lower perivascular space pressures. If the small vessels are covered with endfeet of uniform size across all branching orders, then there are relatively few endfeet and a relatively limited amount of endfoot gaps, resulting in smaller outflows into the interstitial space compared to using the measured small-area endfeet.

Although these comparisons are useful, they are impacted by the fact that the larger vessels have a much larger surface area for flow and solute exchange. It is also helpful to divide the flow by the surface area of the vessel to obtain a flux (i.e. a flow per area). If all the branching orders are covered by uniform area endfeet, then the flux decreases approximately linearly with branching order (**Figure 3E**, pink dots). This is expected since the only variable changing with branching order is the perivascular pressure, which declines approximately linearly with branching order (**Figure 3B**, gray triangles). If different branching orders are covered with endfeet whose areas vary in a manner consistent with the measured values, a wholly distinct pattern is observed. The flow per area (or flux) from the perivascular space into the interstitial is more uniform and independent of branching order, particularly for the middle range of vessel sizes (orders 3-8) over which flux is held nearly constant (**Figure 3E**, purple dots). Compared to values from the model with uniform endfeet, the flux for the largest vessel segments is reduced (by ~25%), while the flux for the smallest vessel segments is markedly increased (>50%). Thus, due to the variance of endfoot size by branching order, flux from the perivascular space into the surrounding interstitium is largely maintained through the cortical depth despite declining perivascular pressure and smaller overall-vessel surface area.

### Perivascular endfeet are larger surrounding arteries than veins

Intracisternally injected tracers enter the cortex primarily along perivascular spaces surrounding penetrating arteries and arterioles rather than ascending veins^12–14^. We next used immunofluorescence double-labeling with a vascular smooth muscle marker (smooth muscle actin, SMA) to evaluate whether the relationship between endfoot dimensions and vessel diameter differed between arterial versus venous segments. Arteries were definitively identified by the pattern of continuous circumferential banding of SMA (**Figure 4A**, top), while veins were identified by the absence of such solid banding, rather exhibiting thin and discontinuous labeling with SMA (**Figure 4A**, bottom). Because differentiation between pre-capillary arteriolar and post-capillary venule segments is difficult, only penetrating or ascending vessels >15 um in diameter that permitted definitive characterization were analyzed. All identifiable vessels from three forebrain coronal sections from each of five wild-type mice were analyzed. Upon first examination, the MLC1 staining surrounding arteries and veins suggested that the endfeet surrounding arteries are distinctly larger compared to similarly-sized veins (**Figure 4A**). For both arterial and venous segments, endfoot size increased with increasing vessel diameter (**Figure 4B**, P_artery_ = 0.0004, R_artery_^2^ = 0.05981. P_vein_ < 0.0001, R_vein_^2^ = 0.08461), although the arteries exhibited a significantly steeper relationship than did veins (Slope_artery_ = 12.18. Slope_vein_ = 3.665. P_artery-vs-vein_ = 0.0052). Similar results were observed when all endfoot measurements were averaged along each individual vessel segment (**Figure 4C**), or when median endfoot dimensions were plotted for each vessel (**Figure 4D**). It is important to highlight that the data used for the modeling study above included data pooled from both arterial and venous segments. These results suggest that the modeled effect of varying endfoot size by vessel branching order (**Figure 3**) likely overestimates the effect along veins, where endfeet remain much more uniformly small along different branching orders, while likely underestimates the effect along arteries, where the influence of branching order on endfoot size is even more pronounced.

**Figure 4:**
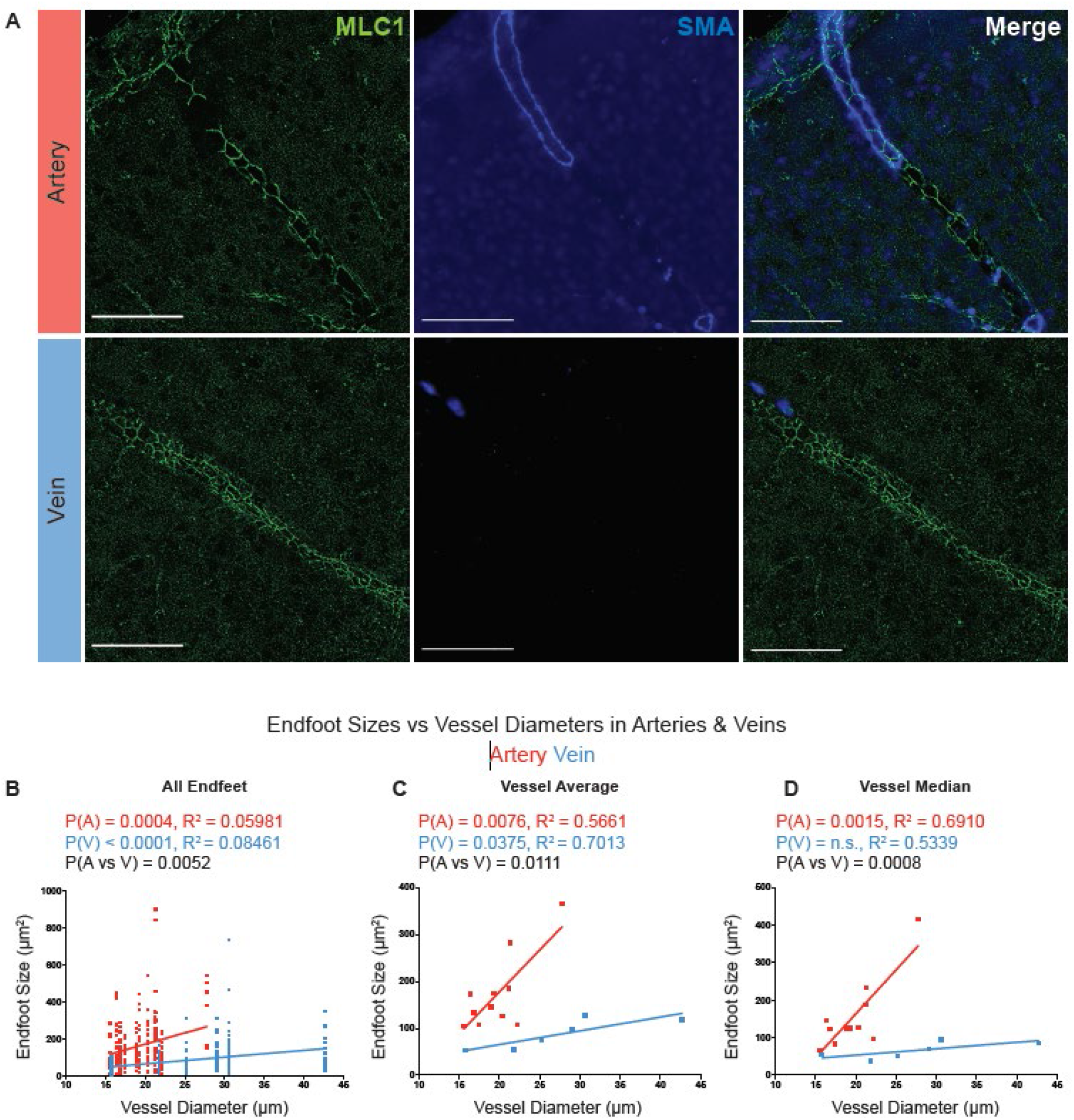
Arteries exhibit greater changes in endfoot size with vessel branching than do veins. **(A)** Immunofluorescence double-labeling with MLC1 and smooth muscle actin (SMA) was used to define endfoot dimensions along arterial (top) and venous (bottom) segments. **(B)** Both arteries and veins exhibited increasing endfoot areas with increasing vessel diameter, however the relationship was significantly (P = 0.0052) steeper among arteries than among veins (P_artery_ = 0.0004, R_artery_^2^ = 0.05981. P_vein_ < 0.0001, R_vein_^2^ = 0.08461). **(C)** Similar analysis using endfoot dimensions averaged for each vessel and median endfoot dimension for each vessel **(D)** showed the same relationships.

## Discussion

In this study, we investigated the anatomy and the biophysical implications of the astrocyte endfoot dimensions surrounding the brain vasculature. We found: (1) The areas of astrocyte endfeet vary by vessel order and diameter; (2) this variation is most pronounced in the arterial, rather than venous vasculature; (3) computational modeling suggests that this heterogeneity of endfeet dimensions maintains relatively constant flux between perivascular and interstitial compartments despite declining pressures along the cerebrovascular tree and through the cortical depth.

### Vessel-dependent astrocyte endfeet heterogeneity

Using MLC1 as a marker of astrocyte endfoot-endfoot junctions along non-capillary vessels, we discovered a heterogeneity in perivascular endfoot dimensions along vessels of different branching orders and diameters. Specifically, the endfeet surrounding large vessels had larger individual surface areas than those surrounding small vessels. The astrocyte endfoot-vessel diameter association was linear, did not differ between young and aged animals, and was more pronounced along arterial versus venous segments. Taken together, these findings provide a novel anatomical insight into astrocyte heterogeneity along the cerebrovascular tree.

Astrocyte heterogeneity has been largely overlooked until recently; in the past ten years increasing evidence demonstrates that astrocytes throughout the central nervous system exhibit substantial morphological, molecular and functional variability. In cortex, protoplasmic astrocytes and fibrous astrocytes have distinct morphologies, localizing predominantly to grey and white matter, respectively^43,44^. Recent studies using single-cell sequencing and in situ techniques reveal a cortical layer- and subcortical region-dependent astrocyte molecular heterogeneity, which suggests that the astrocyte architecture may be regulated by local environmental and neuronal cues^45,46^. Our findings suggest that astrocytes may similarly vary with respect to their association with vessels of different vessel types, different branching order and diameters. This adds another potential layer of astrocyte heterogeneity, although the molecular basis for these differences, and the environmental cues that create and maintain them remain undefined.

Astrocyte processes surround synapses and cover both the relatively open perivascular spaces surrounding penetrating vessels, as well as the basal lamina and extracellular matrix surrounding the intraparenchymal microvasculature. Protoplasmic astrocytes don’t overlap under healthy conditions^47–49^, and the endfeet expansion along the perivascular space surrounding blood vessels is dynamically regulated in injury models^50,51^. It is reasonable to surmise that local cues released from the cerebrovascular endothelial cells, mural cells (vascular smooth muscle cells and pericytes) may affect the sizes of individual endfeet surrounding those vessels, which varies between different vessel types and vessel branching orders. Whether differences in perivascular endfeet dimensions arise developmentally in parallel with the growth of the cerebral vasculature, or are defined locally by the different factors associated with the capillary, arterial and venous micoenvironment is a clear avenue for future inquiry.

### Implications for perivascular glymphatic exchange dynamics

To begin to understand how variation of endfoot dimensions may affect fluid exchange along perivascular spaces, we developed computational models based on two glial-vascular structures: one surrounded by endfeet of uniform dimensions (the mean of the measured endfoot areas), and one surrounded by heterogenous endfeet reflecting the experimentally measured values. The perivascular pressure and flow rate were estimated using the model of Faghih and Sharp ^40^. The magnitude of perivascular pressures has not yet been experimentally defined, and remains an active subject of debate in the field^15,52–54^. It is reasonable, however, to posit that this pressure is impacted by the blood pressure in the vessel that the perivascular space surrounds. The flow rates from the perivascular space and into the interstitial space are normalized and independent of the actual magnitude of perivascular pressure. As anticipated, the vascular tree surrounded by uniform endfeet exhibited a nearly linear drop in normalized flux between the perivascular and interstitial compartment resulting from reduced perivascular pressure along the branching vasculature. In contrast, the vascular tree surrounded by heterogenous endfeet maintained nearly constant perivascular-interstitial flux across seven branching orders. Larger perivascular endfeet along large vessel segments left comparatively fewer endfoot-endfoot junctions (per unit area) for flux, while small endfeet had a greater proportion of endfoot-endfoot junctions contributing to the surface area surrounding small vessels (**Figure 5**). It is possible that this variation in endfoot dimensions along the cerebrovascular tree represents an evolutionary adaptation to maintain efficient exchange between the CSF and interstitial compartments into deeper tissues. Evaluating whether similar heterogeneity is observed in the larger human brain will be an important follow-up study.

**Figure 5:**
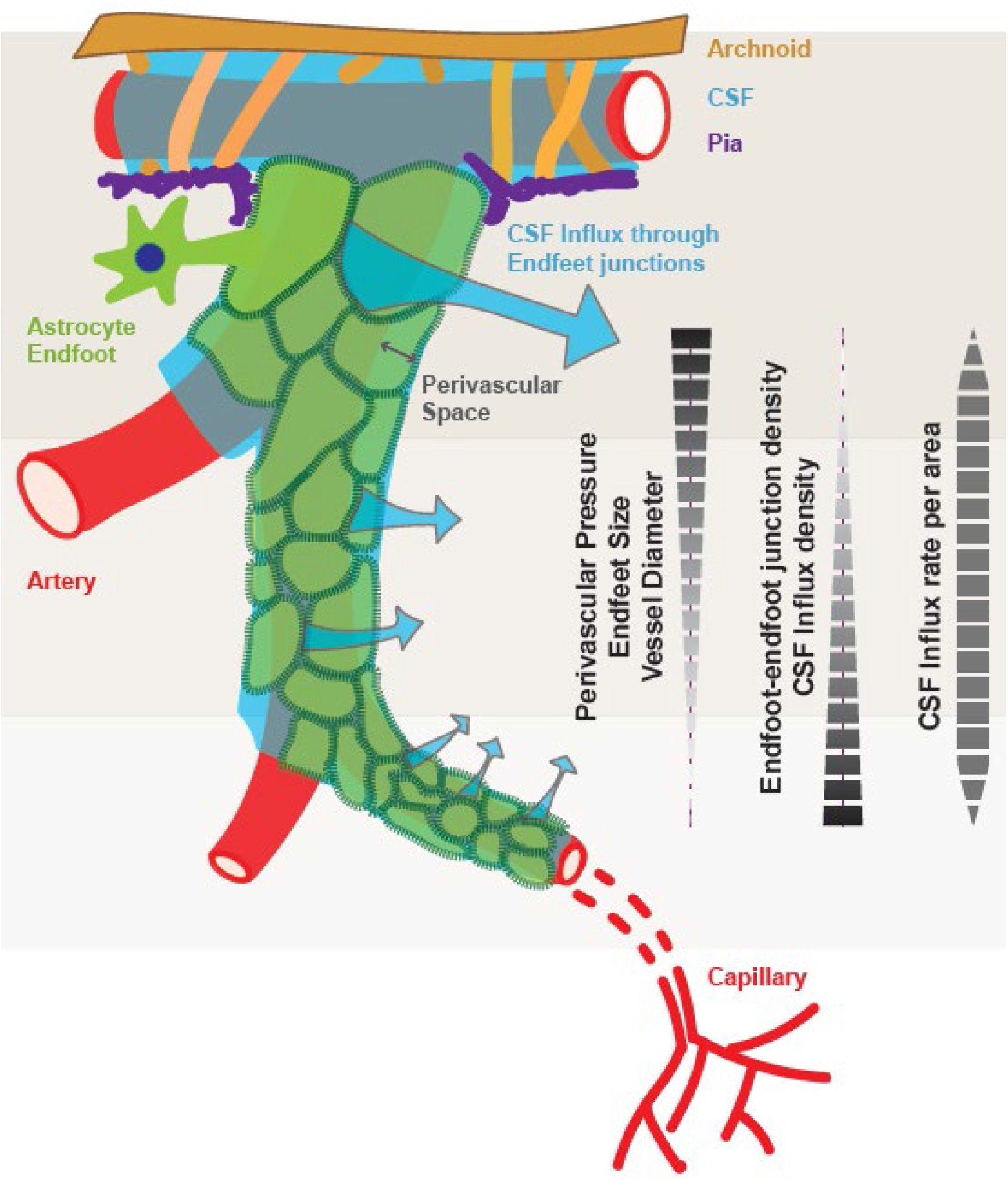
Schematic outline showing the effect of varying endfoot dimensions along the vascular tree on perivascular-interstitial flux. Larger endfoot sizes surrounding large-diameter vessels restrict the relative availability of endfoot-endfoot junctions and reduce superficial perivascular-interstitial flux. As vessels branch, smaller endfoot dimensions increase the relative contribution of endfoot-endfoot junctions to the perivascular boundary, maintaining perivascular-interstitial flux in the face of declining perivascular pressure. The relatively constant perivascular-interstitial flux through several branching orders may reflect an evolutionary adaptation to maintain fluid exchange into deep brain regions far removed from CSF compartments.

The preferential movement of solutes within the CSF and interstitial compartments along peri-arterial spaces has been observed for several decades^11,55,56^. In an early study by Rennels et al.^9^, the protein horseradish peroxidase injected into the cisternal CSF compartment was observed to enter into brain tissue principally along peri-arterial routes. These findings have been confirmed more recently in experimental studies using dynamic imaging approaches such as 2-photon microscopy and dynamic contrast-enhanced magnetic resonance imaging (DCE-MRI) defining the glymphatic system, first in rodents^12–14^ and subsequently in human subjects^38,39^. The basis for this organization of CSF movement along the leptomeningeal and penetrating arterial vasculature has been of substantial interest. Early^55^, and more recent^14,15^ reports suggest that arterial pulsation is a key driver of perivascular fluid movement, and that entrainment of fluid movement along the pulsating arterial vasculature may underlie this organization^57^. The present observation that variation in endfoot dimensions along the cerebrovascular tree is much more pronounced along arteries than along veins suggests that this adaptation may contribute to the preferential movement of CSF solutes into and through brain tissue along perivascular spaces surrounding penetrating arteries, supporting exchange along deeper segments of the vasculature than would be possible with uniform endfoot dimensions. The smaller number of endfoot-endfoot junctions surrounding larger vessels reduce outflow; while greater number of endfoot-endfoot junctions surrounding deeper segments permit increased flux into the interstitial compartment. Indeed, because the computational model was based on pooled data derived from both arterial and venous segments, it is likely that the preservation of perivascular-interstitial flux along arteries is *underestimated* in the present model, while that along veins is *overestimated*.

### Perspectives on endfoot-endfoot junction dynamics and function

The cerebral vasculature is characterized by the presence of tight junctions between endothelial cells^1^. In contrast, the junctions in between the astrocyte endfeet encircling the vasculature have gaps that permit the passage of fluid and macromolecules^58^. Whether surrounding CSF-filled Virchow-Robin spaces encompassing penetrating vessels, or surrounding the basal lamina of the cerebral capillary bed, perivascular astrocytic endfeet cover nearly the entire vascular tree^4,59^. The junctions between astrocyte endfeet serve as the gate for the passage of solutes that are not readily taken up by astroglial membrane channels or transporters into the surrounding interstitial space^58^. However, little is known about the width, dynamics, and regulation of the astrocyte endfoot-endfoot junctions. In this study, the computational model was built based on the assumption that all endfeet gaps had the same width. While this assumption is reasonable due to a lack of experimental data, future studies should seek to define whether these endfoot-endfoot gaps are in fact uniform in their dimensions and what the implications of any non-uniformity may be.

A related unresolved question is whether these endfoot-endfoot gaps remain static, or are rather regulated physiologically. Our modeling results suggest different proportions of endfeet and endfoot-endfoot gaps along the cerebrovascular tree influences perivascular-interstitial flux. Yet if these endfoot junctions themselves were dynamically regulated, potentially through changes in endfoot volume, this might be one way in which perivascular exchange is dynamically regulated. Perivascular endfoot swelling is observed in the setting of osmotic challenge and in the setting of brain injury^60–63^. Astrocytes undergo acute volume regulation in response to osmotic stimuli, a process in which the astroglial water channel AQP4, volume-regulated anion channels, and the transient receptor potential vanillod-4 (TRPV4) cation channel have been variously implicated in^64–66^. One possibility is that dynamic endfoot volume changes, perhaps involving AQP4, regulate perivascular glymphatic exchange. This would provide one potential mechanistic explanation for the observed negative effect of both *Aqp4* gene deletion^12,16^ and loss of perivascular AQP4 localization in the α-syntrophin (*Snta1*) knockout mice^16^ on perivascular glymphatic exchange.

In this study, we used MLC1 as marker of astrocyte endfeet junctions due to its localization to endfoot-endfoot contacts^36^. Although the exact function of MLC1 is unknown, it is suggested to be involved in the regulation of ionic and volume homeostasis through indirect interactions with chloride channel CLC-2^30^, potassium channel Kir4.1^31^, non-selective calcium-permeable channel TRPV4^31^, astrocyte gap junction connexin 43^32^, vacuolar ATPase^33^, and volume activated anion channel (VRAC)^34^. Mutation of *MLC1* gene or GlialCAM gene (*HEPACAM*), an MLC1-interacting protein, causes water accumulation and white matter cysts in the human brain^67^. Therefore, defining the functional role of MLCA MLC1 and its associated proteins in regulating astrocyte endfeet volume and CSF flow dynamics is an intriguing subject for future study.

### Conclusion

Our study described a novel feature of astrocyte heterogeneity, in which perivascular endfoot dimension vary with changes in associated vessel order and diameter. Our computational modeling study defines a physiological consequence of this heterogeneity, in maintaining perivascular-interstitial flux through the cortical depth. These findings suggest that other mechanisms regulating endfoot dynamics may contribute to the physiological regulation glymphatic function.

## Methods

### Animals

Wildtype C57BL/6 mice of 3, 12, and 16-month of age from Jackson Laboratories were housed in cages with free access to food and water and were maintained under controlled day–night cycles in accordance with the NIH Guide for the Care and Use of Laboratory Animals. Mice used in this project were male only. All experiments were approved by the Institutional Animal Care and Use Committee of Oregon Health & Science University.

### Immunochemistry

C57/Bl6 mice were deeply anesthetized with isoflurane and perfused with heparin added saline. Brains were removed and post-fixed in 4% paraformaldehyde in phosphate-buffered saline (PBS) at 4°C overnight. Fixed brains were cut on a Vibratome into 50 μm thickness coronal sections, which were collected in PBS and later transferred to an anti-freezing solution for storage at −20°C. Before immunostaining, sections were rinsed in PBS five times and five minutes each time to wash out any residual anti-freezing solution, mounted and performed antigen retrieval in a citrate buffer (pH = 6) in a steamer for 30 minutes. Both primary and second antibodies were diluted in the same solution as the blocking solution (5% normal donkey serum and 0.3% Triton X-100 in PBS) and incubated overnight. Sections were mounted using Fluoromount-G™ Mounting Medium (ThermoFisher, Cat#: 00-4958-02). Primary antibodies used in this project were anti-MLC1 N-terminal^68^ (guinea pig polyclonal, used at 2 μg/ml), anti-AQP4 (Millipore, Cat#: AB3594, 1:500), Anti-alpha smooth muscle Actin (SMA) (Abcam, ab5694, 1:200). Secondary antibodies used in this project were donkey anti-guinea pig 488 (Jackson, Cat#: 706-545-148, 1:500) and donkey anti-rabbit 594 (ThermoFisher, Cat#: A-21207). DAPI was added with secondary antibody (Abcam, Cat#: ab228549).

### Microcopy and image process

Super resolution z-stack images of the MLC1 and AQP4 co-immunolabeling were acquired on the Zeiss Elyra PS.1. super resolution structured illumination microscope. Images were then deconvolved in the Zeiss imaging acquisition software and reconstructed in image J. Confocal images of the MLC1 and AQP4 co-immunolabeling were acquired on the Zeiss LSM 780 microscope. Widefield images used for quantifying MLC1-labled endfoot sizes were acquired on Zeiss Axioimager, then processed in Image J to subtract background at with a pixel radius of 5.

### Statistics

The linear regression of endfoot areas versus vessel diameters were conducted in Graphpad Prism.

### Model

Fluid flow in the perivascular space is described using an adapted version of the model by Faghih and Sharp ^40^. The model begins by defining a vascular tree with 11 branching orders to span the range of vessel sizes from which endfoot dimensions were collected. The radius of the largest (or root) branch in the vascular tree was set to 23 um, and the radius of daughter branches were approximated using Murray’s law of bifurcations for symmetric trees ^69^:

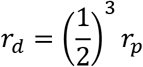

where *r_d_* is the radius of the daughter branch and *r_p_* is the parent-branch radius. The length of each vessel was approximated as 20-times its radius ^70^.

Flow in the perivascular space was assumed to be laminar and the relationship between the flow rate, *Q*, and pressure drop, Δ*P*, for each generation was:

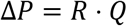

where *R* is the flow resistance. For Poiseuille flow in an annular cross section, the resistance is ^40^:

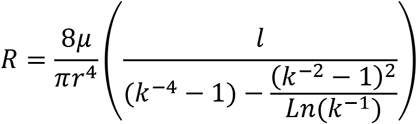

where *k* is the ratio of the inner radius to the outer radius and *μ* is the viscosity (the viscosity of water at body temperature was used). The exact value of *k* is not known, but, importantly, the overall results reported here are normalized so they are independent of the value of *k* used. The total flow rate of fluid in the perivascular space of the model was assumed to be 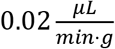 (approximated from ^10,41^), but, again, the presented results are normalized to be independent of this value because it is difficult to accurately measure. In summary, using the approach of Faghih and Sharp ^40^, an idealized, symmetric vascular network is constructed and used to approximate the pressure in the perivascular space across the various generations of the network.

The following steps are taken to map the endfoot-size data onto the idealized vascular network, calculate an average endfoot perimeter for each vessel generation, and simulate the normalized perivascular-to-interstitial flow for each generation:

1. Based on the measured diameter of the actual vessel where each endfoot in the data is located, that same endfoot is assigned to the appropriate generation in the model network.
2. For each generation in the model network, the average endfoot perimeter is calculated by assuming an approximately hexagonal shape. The normalized results are not sensitive to the endfoot shape assumption.
3. For each generation in the model network, the total number of endfeet required to cover the perivascular space is calculated for two cases: (1) the average endfoot size for only the measured endfeet assigned to that generation, and (2) a uniform endfoot size applied to all generations, calculated from the overall average of all endfeet measurements. (This allows comparison between possible outflow based on generationally varying endfoot size and uniform endfoot size across all generations).
4. Based on the number of endfeet for each generation and the average perimeter of the endfeet for that generation, a normalized flow rate for outflow from the perivascular space and into the interstial space is calculated by: *Q_out_* = *L* · *P*/*Q_max_*, where *L* is the total length of endfoot gaps surrounding a vessel (determined in step 3), and *P* is the pressure in the perivascular space (shown in **Figure 3C**). The normalization of this flow rate is based on the largest flow, *Q_max_*, which occurs when the largest branch with the highest pressure in the perivascular space is surrounded by the smaller, uniform endfeet.
5. Finally, because the largest vessels have the largest surface area, the flow rate is divided by the surface area of the vessel to obtain a normalized flow rate per unit surface area (or flux).

